# Rapid cell-free characterization of multi-subunit CRISPR effectors and transposons

**DOI:** 10.1101/2021.10.18.464778

**Authors:** Franziska Wimmer, Ioannis Mougiakos, Frank Englert, Chase L. Beisel

## Abstract

CRISPR-Cas biology and technologies have been largely shaped to-date by the characterization and use of single-effector nucleases. In contrast, multi-subunit effectors dominate natural systems, represent emerging technologies, and were recently associated with RNA-guided DNA transposition. This disconnect stems from the challenge of working with multiple protein subunits *in vitro* and *in vivo*. Here, we apply cell-free transcription-translation (TXTL) to radically accelerate the characterization of multi-subunit CRISPR effectors and transposons. Numerous DNA constructs can be combined in one TXTL reaction, yielding defined biomolecular readouts in hours. Using TXTL, we mined phylogenetically diverse I-E effectors, interrogated extensively self-targeting I-C and I-F systems, and elucidated targeting rules for I-B and I-F CRISPR transposons using only DNA-binding components. We further recapitulated DNA transposition in TXTL, which helped reveal a distinct branch of I-B CRISPR transposons. These capabilities will facilitate the study and exploitation of the broad yet underexplored diversity of CRISPR-Cas systems and transposons.

**HIGHLIGHTS:**

● PAM-DETECT for rapid determination of PAMs for Type I CRISPR-Cas systems in TXTL
● Mining of Type I orthologs and characterization of extensively self-targeting systems
● TXTL-based assessment of DNA target recognition and transposition by CRISPR transposons
● Identification of a distinct branch of Type I-B CRISPR transposons

## INTRODUCTION

CRISPR-Cas systems endow prokaryotes with adaptive defense against invading elements and possess effector nucleases that have become versatile biomolecular tools (Barrangou and Doudna, 2016; Pickar-Oliver and Gersbach, 2019). These systems are remarkably diverse, with two classes, six types, over 30 subtypes, and a few subtype variants defined to-date (Makarova et al., 2019). The two classes are distinguished based on whether the effector nuclease responsible for CRISPR RNA (crRNA)-directed immune defense comprises a multi-protein complex (Class 1) or a single multi-domain protein (Class 2). While systems from both classes have undergone characterization, Class 2 systems have been the most extensively explored. For example, comprehensive determination of target-flanking protospacer-adjacent motifs (PAMs) (Leenay and Beisel, 2017) has been conducted for more than 100 Class 2 effectors spanning at least 15 subtypes (Collias and Beisel, 2021); in contrast, only for 10 Class 1 effectors spanning 7 subtypes (**Table S1**). This discrepancy belies the unique features of Class 1 systems that have attracted increasing attention for basic research and technology development (Hidalgo-Cantabrana and Barrangou, 2020). Class 1 systems represent over 75% of all CRISPR-Cas systems found in nature and contain phylogenetically diverse proteins possessing unique mechanisms of action (Makarova et al., 2015). The associated machinery has also been recently applied as tools in mammalian and plant cells, offering distinct means of achieving programmable gene regulation and genome editing as well as the creation of variable chromosomal deletions (Liu et al., 2018; Zheng et al., 2020). The same machinery has also been associated with emerging alternative functions in bacteria, such as repressing expression of a toxin to promote selection of the CRISPR-Cas system or to counter infection by phages encoding an inhibitory anti-CRISPR protein (Acr) (Li et al., 2021). Finally, a subset of Class 1 systems contain Tn7-like transposon genes and were shown to mediate crRNA-directed transposition (Klompe et al., 2019; Petassi et al., 2020; Peters et al., 2017; Saito et al., 2021). These CRISPR transposons (CASTs) have since been employed in bacteria for the efficient, programmable, and multiplexed insertion of donor DNA exceeding 10 kb (Klompe et al., 2019; Strecker et al., 2019; Vo et al., 2021). The examples noted above highlight the potential of further exploring and harnessing Class 1 CRISPR-Cas systems and CASTs.

The disconnect between the broad relevance of Class 1 systems and the few well-characterized examples can be largely attributed to the challenge of working with multiple protein subunits. Cell-based assays are complicated by the need to encode and optimally express multiple subunits from a minimal number of constructs, while *in vitro* assays require intensive purification of multi-subunit complexes--tasks that are far simpler for single-effector nucleases. A promising alternative came with the advent of cell-free transcription-translation (TXTL) systems and their use for rapidly and scalably characterizing CRISPR-Cas systems (Garamella et al., 2016; Jiao et al., 2021; Liao et al., 2019a, 2019b; Marshall et al., 2018; Maxwell et al., 2018; Silverman et al., 2020; Watters et al., 2018). As part of a TXTL reaction, circular or linear DNA constructs are added to the TXTL mix, resulting in the transcription and translation of the encoded products in minutes to hours. Expressing CRISPR machinery targeted to an included reporter construct further provides a quantitative and dynamic readout based on expression levels and targeting activity. In our prior work, we showed that TXTL could functionally express the Type I effector complex Cascade (CRISPR-associated complex for antiviral defense) that yielded transcriptional repression of a reporter gene (Marshall et al., 2018). However, all other implementations of TXTL to-date have focused on single-effector nucleases (Khakimzhan et al., 2021; Liao et al., 2019a, 2019b; Wandera et al., 2020; Watters et al., 2018). Here, we leverage TXTL to rapidly characterize diverse Type I systems and transposons, allowing ortholog mining, characterization of self-targeting systems, and harnessing of CASTs. The resulting capabilities are expected to accelerate the exploration and exploitation of this broad yet understudied branch of CRISPR biology.

## RESULTS

### PAM-DETECT: a TXTL-based enrichment assay for PAM determination

One of the defining features of DNA-targeting CRISPR-Cas systems is the PAM (Leenay and Beisel, 2017). This collection of sequences always flanks a crRNA target and allows the effector nuclease to discriminate between self (the equivalent targeting spacer in the CRISPR array) and non-self (the invader). However, the associated sequences can vary widely even between close homologs (Collias and Beisel, 2021). Given that the comprehensive PAM determination assays applied for Class 1 systems involved laborious *in vitro* or cell-based assays (**Table S1**), we devised a TXTL-based assay that could elucidate the complete PAM profile recognized by an effector complex but without the need for protein purification or cellular expression (**Figs. 1A and B**). The assay involves expressing the crRNA and the three to five Cas proteins that form Cascade, which then binds target DNA. While Cascade binding normally recruits the endonuclease Cas3 to nick and processively degrade the non-target strand of DNA (Huo et al., 2014; Mulepati and Bailey, 2013; Westra et al., 2012), Cascade strongly binds DNA even without Cas3 (Jore et al., 2011; Westra et al., 2012). As part of the TXTL-based assay, Cascade binds target DNA flanked by a library of potential PAM sequences. After sufficient time to produce Cascade and ensure DNA binding, a restriction enzyme is introduced that cleaves a sequence within the DNA target. As a result, DNA containing a recognized PAM sequence is protected by the bound Cascade, thereby enriching this sequence within the library. Next-generation sequencing (NGS) is then performed to quantify the relative frequency of each PAM sequence before and after restriction digestion. We call this assay PAM-DETECT (PAM-DETermination with Enrichment-based Cell-free TXTL). From the addition of the DNA constructs to the isolation of library DNA for NGS, the entire process requires 13 to 23 hours -- substantially faster than the days to weeks required for *in vitro* and cell-based assays when starting with DNA expression constructs. Also, because the reactions are conducted in a few microliters, reactions can be conducted in parallel in microtiter plates for characterizing a massive number of systems and conditions at one time.

**Figure 1.**
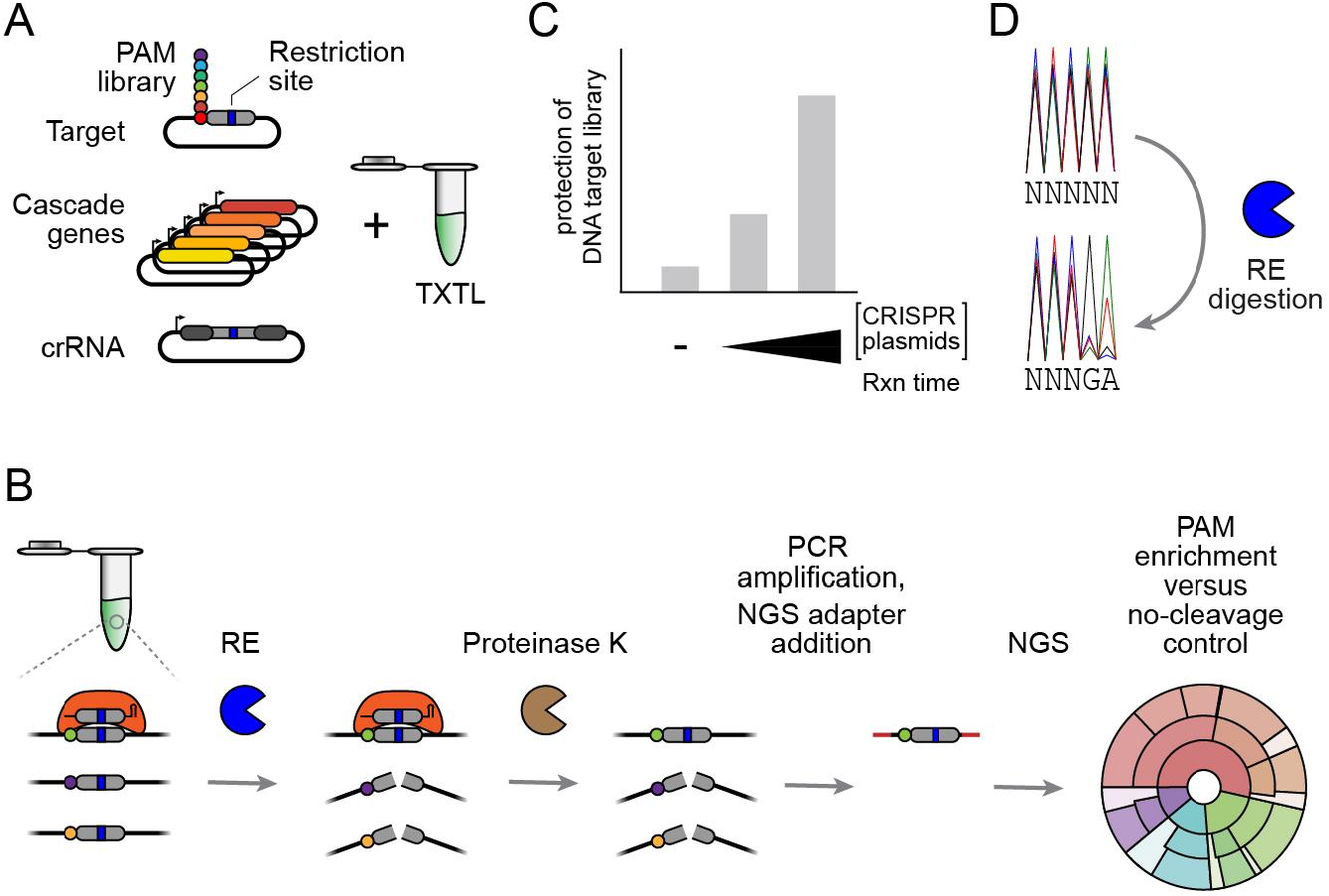
PAM-DETECT, a TXTL-based PAM determination assay for multi-protein CRISPR effectors. (**A**) DNA components added to a TXTL reaction to perform PAM-DETECT. The Cascade genes can be encoded on separate plasmids as shown here or as an operon. (**B**) Steps comprising PAM-DETECT. RE: restriction enzyme. (**C**) Determination of library protection from restriction cleavage by qPCR. A reaction conducted without the Cascade and crRNA plasmids serves as a negative control. (**D**) Determination of PAM enrichment by Sanger sequencing.

As part of PAM-DETECT, we devised two parallel checkpoints to assess the extent of library protection and PAM enrichment prior to submitting samples for NGS. For the first checkpoint (**Fig. 1C**), qPCR is applied with a digested and undigested library to measure the extent to which the library was protected by Cascade binding. Given that excess effector can boost the prevalence of less-preferred PAM sequences (Karvelis et al., 2015), the qPCR results can indicate the stringency of the determined PAM sequences. Fortunately, the conditions of PAM-DETECT can be readily tuned by changing the concentration of the added DNA constructs and the time allowed for Cascade expression and DNA binding. For the second checkpoint (**Fig. 1D**), the digested and undigested libraries are subjected to Sanger sequencing. Elevated peaks in the digested sample reflect enrichment of those bases at that PAM position, providing an early indication of the determined PAM.

### PAM-DETECT validated with the canonical Type I-E CRISPR-Cas system from *Escherichia coli*

To evaluate PAM-DETECT, we began with Cascade encoded by the Type I-E CRISPR-Cas system from *Escherichia coli* (**Fig. 2A**), the best studied Type I system to-date. As part of its extensive characterization, the effector complex has been subjected to multiple comprehensive PAM determination assays (Caliando and Voigt, 2015; Fineran et al., 2014; Fu et al., 2017; Leenay et al., 2016; Musharova et al., 2019; Xue et al., 2015), establishing a complex landscape principally composed of the canonical PAM sequences AAG, AGG, ATG, and GAG (written 5′ to 3′) located on the non-target strand immediately upstream of the guide sequence. We applied PAM-DETECT by encoding the five Cascade genes and a targeting single-spacer CRISPR array encoding a crRNA on six separate plasmids and combining these plasmids with a 5-base PAM target library in TXTL (**Fig. 2A**). To explicitly evaluate the impact of excess effector complexes, we tested two different conditions: one with 0.25 nM of Cascade-encoding plasmids and 6-hour reaction time for low Cascade expression and binding, and another with 3 nM of Cascade-encoding plasmids and 16-hour reaction time for high Cascade expression and binding. The intermediate qPCR check showed significant DNA protection compared to the control lacking Cascade, with ∼2-fold more protection with the high versus low Cascade condition (**Fig. 2B**). Correspondingly, the Sanger sequencing checkpoint showed enrichment of an AAG motif compared to the undigested control, where the motif was more pronounced for the low Cascade condition (**Fig. 2C**). The checkpoints were in line with protection of DNA sequences related to the known PAM, with enhanced protection for the high Cascade condition.

**Figure 2.**
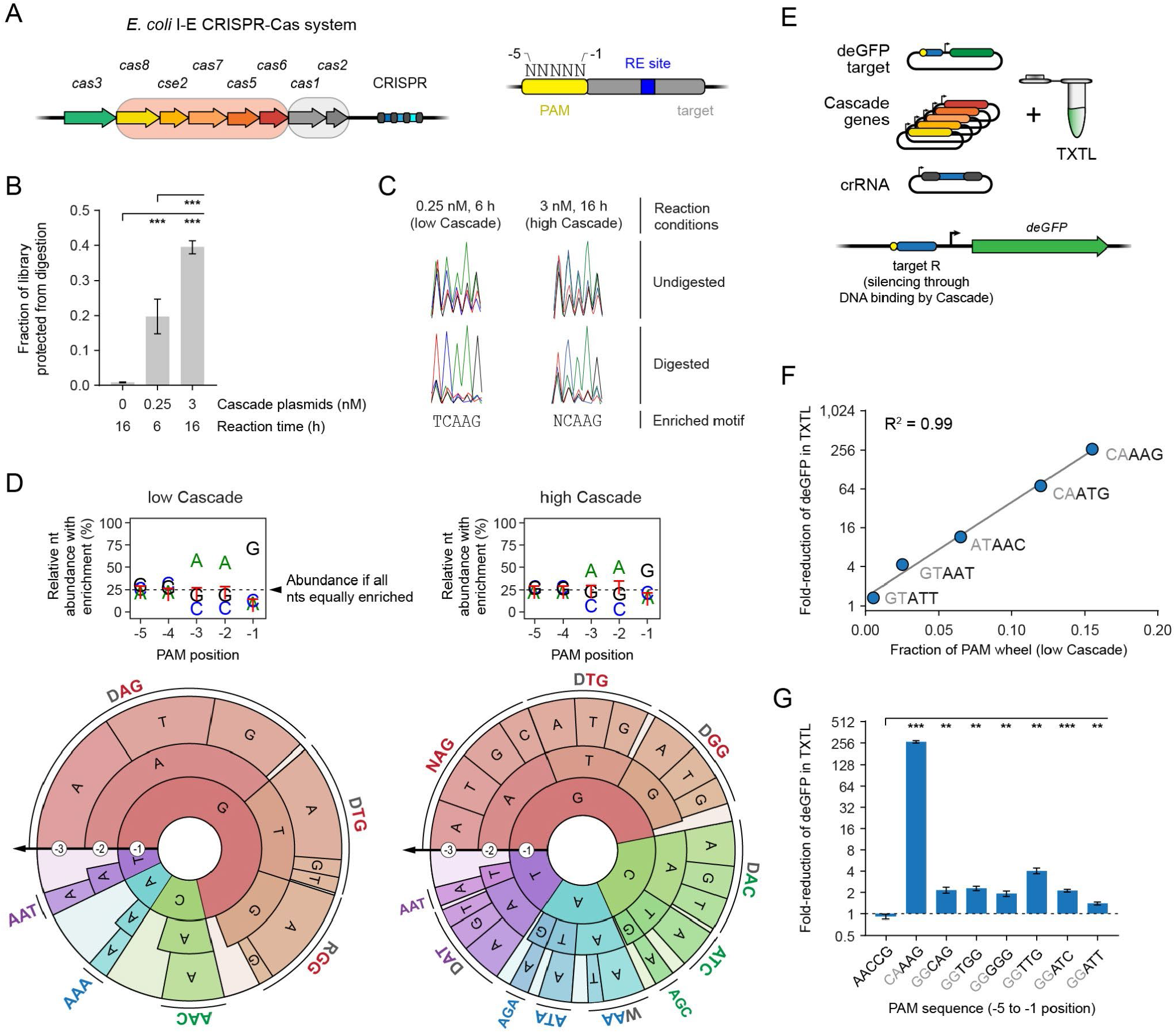
Validation of PAM-DETECT with the I-E CRISPR-Cas system from *E. coli*. (**A**) The Type I-E CRISPR-Cas systems from *E. coli*. The genes encoding the Cascade complex are in the light orange box, while the genes encoding the acquisition proteins are in the gray box. Right: 5N library of potential PAM sequences used with PAM-DETECT. (**B**) Extent of PAM library protection under conditions resulting in low or high levels of Cascade based on qPCR. Library protection compares the library with and without RE digestion. (**C**) Preliminary recognized PAM with low or high levels of Cascade based on Sanger sequencing. Overrepresentation of T and C at the -5 and -4 position, respectively, can be explained by the library generation, as TCAAG represented the most prevalent sequence in the library. As a result, protection of an AAG motive protects the majority of the TCAAG sequences. (**D**) Nucleotide-enrichment plots and PAM wheels based on conducting PAM-DETECT with low or high levels of Cascade. Individual sequences comprising at least 2% of the PAM wheel are shown. Results represent the average of duplicate independent experiments. The size of the arc for an individual sequence corresponds to its relative enrichment within the library. (**E**) Overview of the TXTL-based PAM validation assay. PAM sequences are tested by Cascade binding target R flanked by the tested PAM. Because target R overlaps the promoter driving expression of deGFP, target binding would block deGFP expression. Target R is distinct from the restriction site-containing target used with PAM-DETECT. (**F**) Correlation between PAM enrichment from PAM-DETECT and gene repression in TXTL. Enrichment was based on the fraction of the PAM wheel derived from the low Cascade condition. Enrichment values represent the mean of duplicate PAM-DETECT assays, while fold-reduction values represent the mean of triplicate TXTL assays. Fold-reduction was calculated based on a non-targeting crRNA control. (**G**) TXTL validation of PAM sequences identified by PAM-DETECT but not by PAM-SCANR. CAAAG serves as a positive control. AACCG matches the 3′ end of the repeat and therefore serves as a negative control. The AACCG self PAM is the reference for statistical analyses. Error bars in B and G indicate the mean and standard deviation of triplicate independent experiments. ***: p < 0.001. **: p < 0.01. *: p < 0.05. ns: p > 0.05.

Given the promising results from the two checkpoints, we proceeded to NGS with both Cascade conditions to map the full PAM profile. After determining an enrichment score for each library sequence, we visualized the results as a PAM wheel to capture both individual sequences and enrichment scores (Leenay et al., 2016) (**Fig. 2D**). The PAM wheel for the low Cascade condition captured the four known canonical PAMs as well as other well-recognized PAM sequences (e.g. TAG, AAC) reported in prior screens (Caliando and Voigt, 2015; Fineran et al., 2014; Leenay et al., 2016; Musharova et al., 2019; Xue et al., 2015). The PAM wheel for the high Cascade condition included these PAM sequences as well as other PAM sequences that were less enriched (e.g. AAA, AAT) or negligibly enriched (e.g. CAG, ATT) for the low Cascade condition (**Fig. 2D**). The differences in PAM profiles demonstrate how PAM-DETECT can be readily tuned by varying plasmid concentration and reaction time.

To validate the results, we applied TXTL to silence expression of a deGFP reporter (Shin and Noireaux, 2012) using a distinct target sequence overlapping the reporter’s upstream promoter (**Fig. 2E, Table S2**). The PAM region could then be altered without affecting the promoter sequence. For representative PAM sequences, the fold-repression of deGFP production versus a non-targeting control strongly correlated with the enrichment score of each sequence in PAM-DETECT for the low Cascade condition (R^2^ = 0.99) (**Fig. 2F**). The correlation was particularly striking given the use of a different target sequence, which can affect the apparent hierarchy of PAM recognition (Leenay et al., 2016; Xue et al., 2015). Applying the same assay to PAM sequences enriched under the high Cascade condition but not detected with our previous PAM-SCANR method (Leenay et al., 2016), we measured modest but significant deGFP repression (**Fig. 2G**). These validation experiments show that PAM-DETECT can produce comprehensive and quantitative PAM profiles, and the assay conditions can be readily altered to tune the stringency of PAM detection.

### Distinct PAM profiles pervade I-E CRISPR-Cas systems

After validating PAM-DETECT using the established I-E system from *E. coli*, we turned to the first important use of this assay: mining diverse CRISPR effector proteins and complexes. Nuclease mining has been highly successful for single-effector nucleases such as Cas9, which revealed a wide collection of nucleases recognizing the full spectrum of PAMs (Gasiunas et al., 2020; Zetsche et al., 2020). Nuclease mining therefore could be highly valuable when applied to Class 1 systems. Focusing again on the I-E subtype of CRISPR-Cas systems, we began by identifying diverse Cas8e proteins responsible for PAM recognition within Cascade from known cultured mesophilic bacterial strains. This analysis revealed a set of 213 Cas8e proteins (**Table S3**). We further divided the Cas8e set in groups according to the amino-acid sequence of the highly variable L1 loop within the N-terminal domain (**Table S3**) reported to stabilize the Cas8e-PAM interactions (Tay et al., 2015; Xiao et al., 2017). The numerous clusters with distinct L1 motifs suggested diverse modes of PAM recognition extending beyond that observed with *E. coli*’s Cascade.

We selected 11 representative I-E systems reflecting some of the most abundant L1 motifs to characterize with PAM-DETECT (**Figs. 3A, S1**). Characterizing the resulting Cascade complexes required encoding 55 Cascade genes and 11 single-spacer arrays, each in separate plasmids. However, despite this large number of constructs, PAM-DETECT could be performed with all constructs in parallel. We selected the high Cascade conditions (3 nM plasmids, 16 hour reaction time) given uncertainty about how well a given system would be functionally expressed in TXTL. All but one system yielded significant enrichment of the PAM library compared to a non-digested control (**Fig. S1A**), allowing us to determine a large number of PAM profiles.

**Figure 3.**
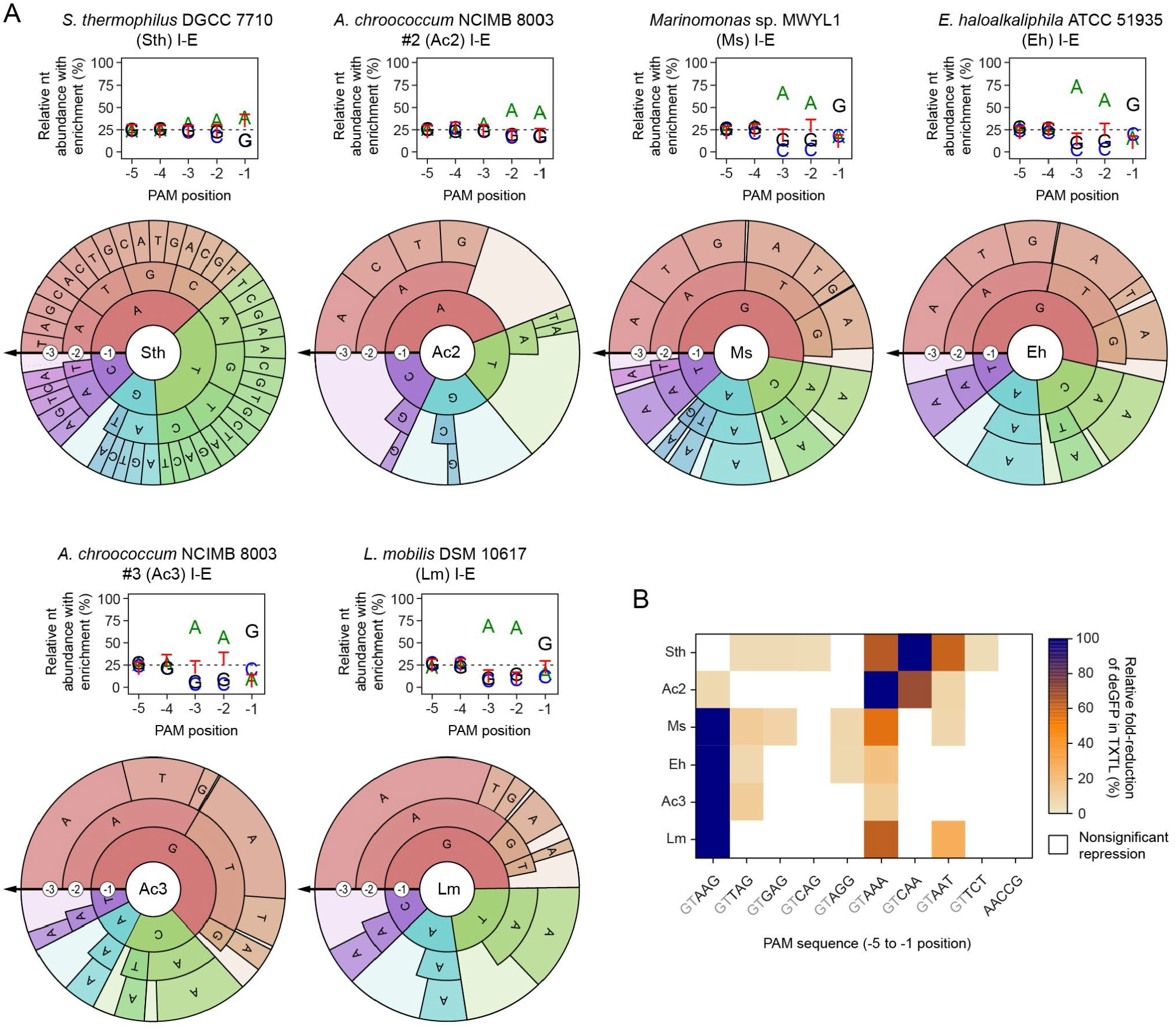
Harnessing the functional diversity of I-E CRISPR-Cas systems. (**A**) Nucleotide enrichment plots and PAM wheels for selected I-E systems subjected to PAM-DETECT. See Figure S1 for 5 additional systems subjected to PAM-DETECT. Ac1 (in Figure S1), Ac2, and Ac3 are present in the same bacterium. Individual sequences comprising at least 2% of the PAM wheel are shown. Plots and PAM wheels are averages of duplicate independent experiments. (**B**) Comparison of PAM recognition between systems. Recognition was determined by assessing repression of a deGFP reporter in TXTL. Values represent the mean of three TXTL experiments. Fold-reduction values that are not significantly different from that of the non-targeting crRNA control (p > 0.05) are shown as white squares. The PAM sequence showing the highest fold-reduction for each system was set to 100%. AACCG matches the 3′ end of the repeat for most of the systems.

PAM-DETECT revealed a broad range of recognized PAMs (**Figs. 3A, S1B**). The PAM profile most distinct from that associated with the *E. coli* Cascade was recognized by Cascade from *Streptococcus thermophilus* DGCC 7710 (Sth), which recognized any sequence with an A or T at the -1 position as well as AS (S = G, C) and ATS. While the *S. thermophilus* Cascade protected ∼75% of the library -- indicative of enriched sub-optimal PAMs, the PAM profile matched the few PAM sequences previously confirmed to bind purified Cascade *in vitro* (Sinkunas et al., 2013). Most remaining systems generally recognized AAG as a dominant PAM sequence, although there were notable deviations and additions. For example, one system from *Azotobacter chroococcum* NCIMB 8003 (Ac2) principally recognized AA, while another system from *Paracoccus* sp. J4 (Ps) preferentially recognized AAC. Separately, the systems from *Marinomonas* sp. MWYL1 (Ms), and *Ectothiorhodospira haloalkaliphila* ATCC 51935 (Eh) as well as a separate system in *Azotobacter chroococcum* NCIMB 8003 (Ac3) recognized PAM profiles paralleling that recognized by *E. coli*’s system. Notably, Ac2 and Ac3 are present in the same bacterium, suggesting that their partially overlapping PAM profiles could confer redundancy in immune defense as reported for co-occurring Type I and Type III systems (Silas et al., 2017). The distinct PAM profiles that gave measurable activity in the deGFP silencing assay in TXTL confirmed the trends observed with the PAM wheels (**Figs. 3B**). Given that Type I-E systems represent one of the most abundant CRISPR-Cas subtypes in nature (Makarova et al., 2015), our initial characterization suggests that a far greater diversity of recognized PAM profiles likely exists across this expansive subtype.

### Extensively self-targeting I-C and I-F1 CRISPR-Cas systems in *Xanthomonas albilineans* are functionally encoded

Beyond mining individual systems, PAM-DETECT can be further applied to interrogate systems that deviate from traditional immune defense. Prominent examples are self-targeting CRISPR-Cas systems that encode crRNAs targeting chromosomal locations (Wimmer and Beisel, 2019). While self-targeting is considered inherently incompatible with a functional CRISPR-Cas system (Gomaa et al., 2014; Stern et al., 2010; Vercoe et al., 2013), accumulating examples provide important counterpoints where the systems tolerate or even utilize self-targeting crRNAs. For instance, systems encoding self-targeting crRNAs have been associated with prophage-encoded Acrs that actively repress immune defense and serve as markers to uncover novel Acrs (Marino et al., 2018; Rauch et al., 2017; Watters et al., 2018; Yin et al., 2019). Furthermore, a crRNA-like RNA encoded within Type I systems was also shown to direct Cascade to a partially complementary site upstream of a toxin gene, thereby blocking its transcription to ensure maintenance of the CRISPR-Cas system and counter Acr-encoding phages (Li et al., 2021). PAM-DETECT and TXTL therefore could accelerate the characterization of these unique systems.

We specifically focused on two extensively self-targeting CRISPR-Cas systems within the plant pathogen *Xanthomonas albilineans* CFBP7063. This bacterium encodes two CRISPR-Cas systems (I-C and I-F1) each harboring the full cohort of *cas* genes and associated with a remarkably large repertoire of self-targeting spacers (**Fig. 4A**). Of the 64 spacers present across the six CRISPR arrays, 24 (38%) at least partially match sites in the chromosome or one plasmid (**Table S4, Fig. S2A**) with a common set of flanking PAMs (**Fig. 4B**). TXTL therefore offered a rapid means to explore the functionality of these systems and why self-targeting is tolerated.

**Figure 4.**
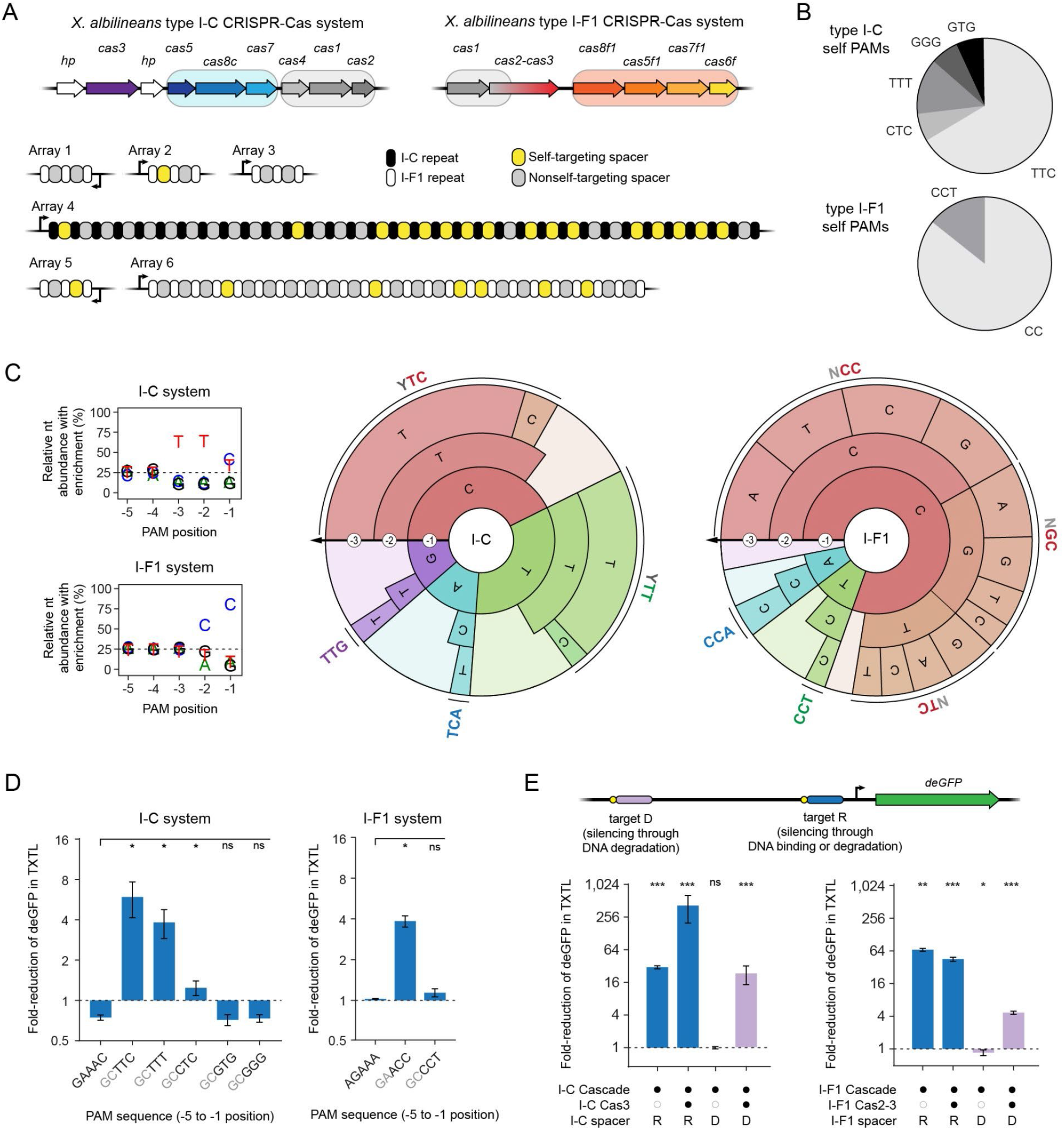
Interrogating extensive self-targeting for two type I CRISPR-Cas systems in *Xanthomonas albilineans*. (**A**) Overview of the I-C and I-F1 CRISPR-Cas systems and self-targeting spacers. The genes encoding the Cascade complex are in the light blue box (I-C) or the light orange box (I-F1), while the genes encoding the acquisition proteins are in the gray box. (**B**) Distribution of PAMs associated with the self-targets. See Figure S2 for the self-target location and Table S4 for the self-target sequences. (**C**) Nucleotide-enrichment plots and PAM wheels based on conducting PAM-DETECT. Individual sequences comprising at least 2% of the PAM wheel are shown. Plots and PAM wheels are averages of duplicate independent experiments. (**D**) Validation of PAMs associated with self-targets in TXTL. Fold-reduction was calculated based on a non-targeting crRNA control. GAAAC and AGAAA match the 3′ end of the repeat for the I-C and I-F1 systems, respectively. Either self PAM is the reference for statistical analyses. (**E**) Assessing DNA binding by Cascade and DNA degradation by Cas3 in TXTL. Targeting far upstream of the promoter (target D) can reduce deGFP levels only through degradation of the plasmid. Targeting the promoter (target R) can reduce deGFP levels through DNA binding or plasmid degradation. Fold-reduction was calculated based on a non-targeting crRNA control. The non-targeting crRNA control is the reference for statistical analyses. Target D with only the I-F1 Cascade yielded modestly but significantly altered deGFP levels between targeting and non-targeting conditions, although targeting resulted in an increase in deGFP levels. Errors bars in D and E indicate the mean and standard deviation of triplicate independent experiments. ***: p < 0.001. **: p < 0.01. *: p < 0.05. ns: p > 0.05.

We first performed PAM-DETECT using Cascade from both CRISPR-Cas systems (**Fig. 4C**). Either Cascade protected a small portion of the DNA library (∼2% for I-C, ∼6% for I-F1) from restriction digestion (**Fig. S2B**), indicating functional expression of all Cascade subunits. PAM-DETECT further revealed PAM profiles that overlapped -- but were not identical to -- the I-C and I-F1 systems with even a moderately mapped PAM profile (Almendros et al., 2012; Leenay et al., 2016; Rao et al., 2017; Rollins et al., 2015; Tuminauskaite et al., 2020; Zheng et al., 2019). In particular, the I-C system from *X. albilineans* recognizes TTC followed by TTT and CTC, while the characterized I-C system from *Bacillus halodurans* recognizes TTC followed by CTC and then TCC (Leenay et al., 2016) and the I-C system from *Legionella pneumophila* recognized TTC followed by TTT and CTT (Rao et al., 2017). Separately, the I-F1 system from *X. albilineans* recognizes CC as the strongest PAM similar to other I-F systems (Almendros et al., 2012; Rollins et al., 2015; Tuminauskaite et al., 2020; Zheng et al., 2019), although *X. albilineans* system also can recognize a G and T but not an A at the -2 position and could tolerate a CC PAM shifted one nucleotide upstream. The recognized PAMs of both I-C and I-F1 systems further overlapped with the PAM sequences flanking the self-targets for 87% of the I-C self-targets (TTC, TTT, CTC) and all I-F1 self-targets (CC, CCT) (**Figs. 4B** and **C**). Testing these individual PAMs in TXTL using gene repression with Cascade confirmed that the I-C system could recognize not only TTC but also TTT and CTC (**Fig. 4D**). The same TXTL assay confirmed that the I-F1 system could recognize the CC PAM associated with almost all self-targeting. PAM-DETECT therefore can be implemented beyond I-E systems and indicated that the interrogated I-C and I-F1 systems in *X. albilineans* are capable of binding the vast majority of self-targeting sites in the genome.

If the Cas3 endonuclease for either system is functionally encoded and expressed, then recognition of these self-targeting sites should prove lethal to this bacterium. We therefore reconfigured the TXTL assay to evaluate the extent to which the I-C or I-F1 Cas3 could elicit DNA degradation (**Fig. 4E**). The DNA target was placed in the backbone of the deGFP reporter ∼200 bps upstream of the deGFP promoter flanked by a TTC (I-C) or CC (I-F1) PAM, which would only lead to loss of deGFP fluorescence if the backbone is nicked or cleaved, leading to DNA degradation by RecBCD (Marshall et al., 2018). For both systems, this new target site location resulted in targeted deGFP silencing following expression of Cascade and Cas3 but not Cascade alone (**Fig. 4E**). Cas3 is therefore functionally encoded and would lead to lethal self-targeting unless Cascade is fully silenced in this bacterium or another mechanism is in place to inhibit Cascade and/or Cas3 activity. The findings thus lay a foundation to investigate the mechanistic basis of self-targeting and whether self-targeting underlies functions extending beyond immune defense.

### The I-F CRISPR transposon from *Vibrio cholerae* recognizes an extremely flexible PAM profile

The demonstrated applicability of PAM-DETECT for diverse Type I CRISPR-Cas systems created a unique opportunity: applying the same assay to CASTs. Of the three known CAST types (I-B, I-F, V-K), two (I-B, I-F) rely on Cascade for DNA target recognition (Klompe et al., 2019; Saito et al., 2021). Recognition then leads to integration of the transposon DNA at a defined distance downstream of the target. Characterization of these systems to-date has relied on encoding a crRNA, all CRISPR and transposon components, and donor DNA flanked by the transposon ends in bacteria to achieve targeted transposition. However, the reliance of I-B and I-F CASTs on Cascade offers an opportunity to express only these CAST components as part of PAM-DETECT to elucidate key rules for DNA target recognition.

We began with the I-F CAST from *V. cholerae* that exhibited robust DNA integration in *E. coli* and has been used for multiple applications in bacteria (Klompe et al., 2019; Vo et al., 2021) (**Fig. 5A**). Prior screening of individual potential PAM sequences via transposition in *E. coli* revealed a general preference for a C at the -2 position, although a comprehensive PAM remained to be determined. We therefore applied PAM-DETECT by expressing the three Cascade genes (a natural *cas8-cas5* fusion, *cas6*, and *cas7*) along with the *tniQ* gene responsible for recruiting the other three transposon genes (*tnsA*, *tnsB*, *tnsC*), as the role of TniQ in DNA target recognition remained to be established (Klompe et al., 2019; Petassi et al., 2020; Vo et al., 2021). PAM-DETECT revealed 57% DNA protection under high Cascade conditions (3 nM plasmids, 16 hour reaction time), leading us to also perform PAM-DETECT with the low Cascade conditions (0.25 nM plasmids, 6 hour reaction time) that exhibited 25% DNA protection (**Fig. S3A**). We further found that *tniQ* was dispensable for DNA binding (**Fig. S3B**). The resulting PAM profile was remarkably flexible, with a preference for a C and bias against an A at the -2 position (**Figs. 5B, S3C**). We further noticed deviations from these biases that could still allow target recognition. For example, recognition of a G or T at the -2 position could be enhanced with a C at the -1 position or an A at the -3 position. Separately, an A at the -2 position could be rescued with a C at the -3 position (**Figs. 5B, S3C**). The results from PAM-DETECT therefore suggest that this I-F CAST recognizes a remarkably flexible PAM profile with preferences extending beyond a simple consensus sequence.

**Figure 5.**
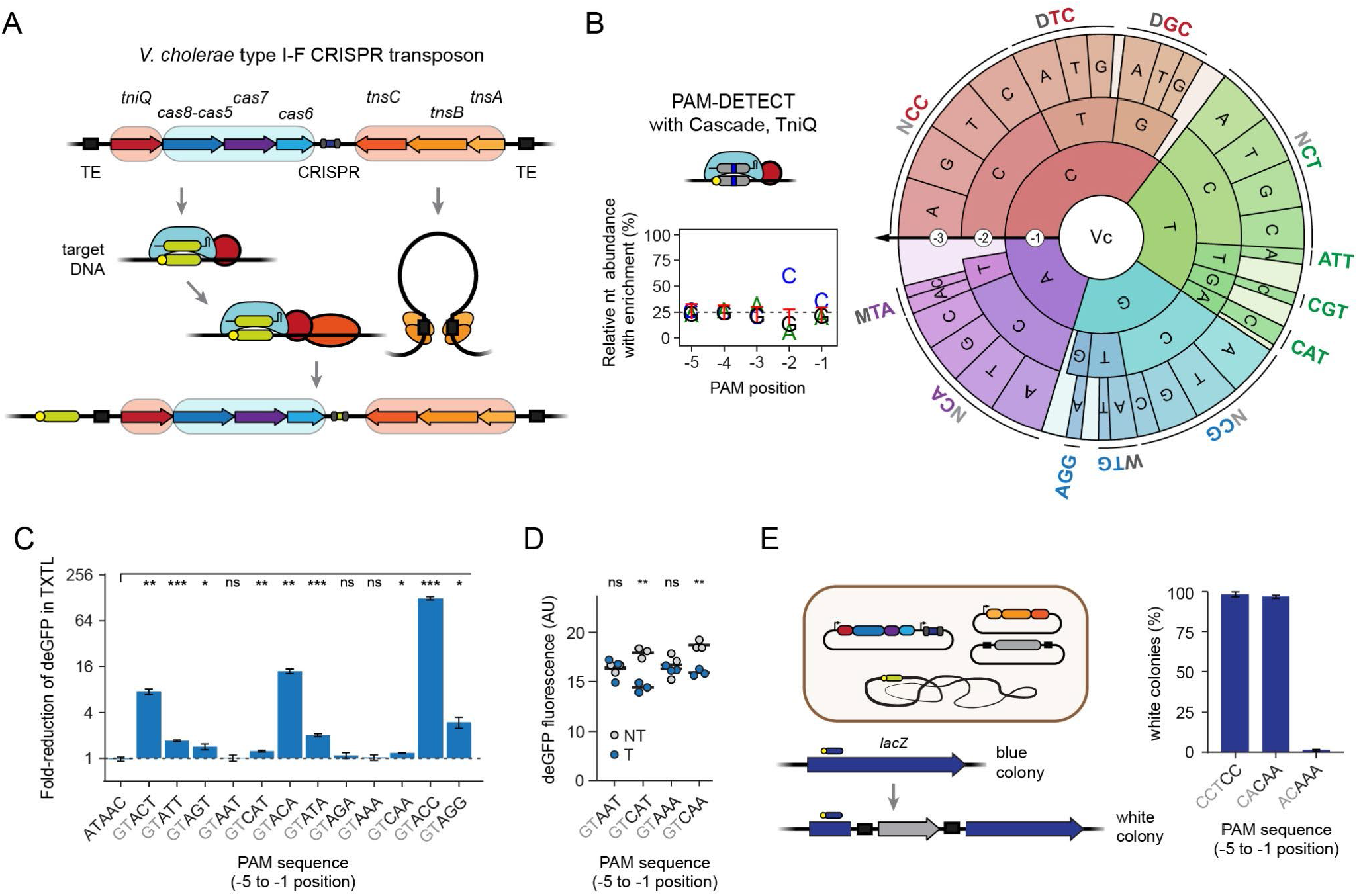
Interrogating the PAM profile of the *Vibrio cholerae* I-F CRISPR transposon. (**A**) Overview of *V. cholerae* I-F CRISPR transposon and its mechanism of transposition. (**B**) Nucleotide-enrichment plot and PAM wheel based on conducting PAM-DETECT with Cascade and TniQ. Individual sequences comprising at least 1% of the PAM wheel are shown. The plot and PAM wheel are averages of duplicate independent experiments. (**C**) Validation of PAMs in TXTL. Gene repression was evaluated with Cascade and the indicated PAM flanking target R upstream of the deGFP reporter. See Figure 2E for details. Fold-reduction was calculated based on a non-targeting crRNA control. ATAAC matches the 3′ end of the repeat and therefore serves as a negative control. The ATAAC self PAM is the reference for statistical analyses. (**D**) Individual measurements of endpoint deGFP levels in TXTL. Triplicate values are shown for selected PAMs with a targeting (T) or non-targeting (NT) crRNA. See C for details. (**E**) Validation of PAM recognition for DNA transposition in *E. coli*. Donor DNA is inserted within the *lacZ* gene, preventing the formation of blue colonies on IPTG and X-gal. Different targets within *lacZ* were selected to test the indicated PAM. The targets for the CAA and AAA PAMs are shifted by one nucleotide. See Figure S3 for more information. Error bars in C, D, and E indicate the mean and standard deviation of triplicate independent experiments. ***: p < 0.001. **: p < 0.01. *: p < 0.05. ns: p > 0.05.

To evaluate the PAM profile output by PAM-DETECT, we first employed our TXTL-based deGFP silencing assay (**Figs. 5C**). Cascade most strongly recognized PAM sequences with C at the -2 position, with the greatest preference for CC. Deviating from this preference reduced but did not eliminate measurable silencing as long as A was not present at the -2 and -3 positions. Interestingly, while AAA and AAT yielded no measurable deGFP silencing, replacing A with C at the -3 position restored measurable silencing, albeit with low activity (**Fig. 5D**). These small but measurable differences raised the question of how these activities translate into programmable DNA transposition in *E. coli*. We therefore employed the previously described transposition system in which the CAST genes and crRNA are encoded outside of donor DNA flanked by the transposition ends (Klompe et al., 2019), and transposition is conducted at 30°C for higher integration efficiency (Vo et al., 2021). The crRNA is further designed to drive transposition into the *lacZ* gene in the *E. coli* genome, which yields white rather than blue colonies on the cleavable dye X-gal. Using this experimental setup, we found that a CAA but not AAA PAM sequence yielded robust DNA transposition, even though the targets were separated by only one base (**Figs. 5E, S3D and E**). Furthermore, the measured transposition efficiency was similar for CAA and CC. Therefore, even low levels of gene silencing with Cascade in TXTL could yield efficient transposition in *E. coli*.

### The I-B2 CRISPR transposon from *Rippkaea orientalis* recognizes a less flexible PAM profile

Building on our success applying PAM-DETECT to the I-F CAST from *V. cholerae*, we turned to I-B CASTs. Two examples of I-B CASTs were experimentally characterized very recently, revealing that a second encoded *tniQ* (renamed *tnsD*) drives DNA transposition at conserved sites flanking tRNAs or *glmS* independently of Cascade or a crRNA (Saito et al., 2021). These examples were also previously subjected to a high-throughput PAM determination assay conducted by performing transposition *in vivo* expressing all components in *E. coli*. Type I-B CASTs were further split into two subtypes (I-B1, I-B2) based on the TnsA and TnsB being fused or separate proteins, the general genetic organization of the CAST locus, and crRNA-independent insertion flanking tRNAs or *glmS*.

While exploring examples within the I-B CASTs, we noticed a further division within the I-B2 subtype typified by *tnsD* flanking the Cascade genes rather than the other transposon genes (**Fig. 6A**). This organization more closely paralleled that of I-B1 CASTs (Saito et al., 2021) but still possesses the *tnsAB* fusion and the presence of tRNAs flanking the CASTs indicative of I-B2 CASTs. The division of the I-B2 CASTs in two clades, denoted hereafter as I-B2.1 and I-B2.2, was further supported by the higher shared similarity of the TnsAB, TnsC, TnsD and TniQ proteins from systems that belong to each clade (**Figs. 6A, S4A**). The Cascade proteins were similar across all I-B CASTs and thus could not help differentiate any divisions within this CAST type (Saito et al., 2021). We chose the I-B2.2 CAST from *Rippkaea orientalis* (RoCAST) as a representative example to characterize.

**Figure 6:**
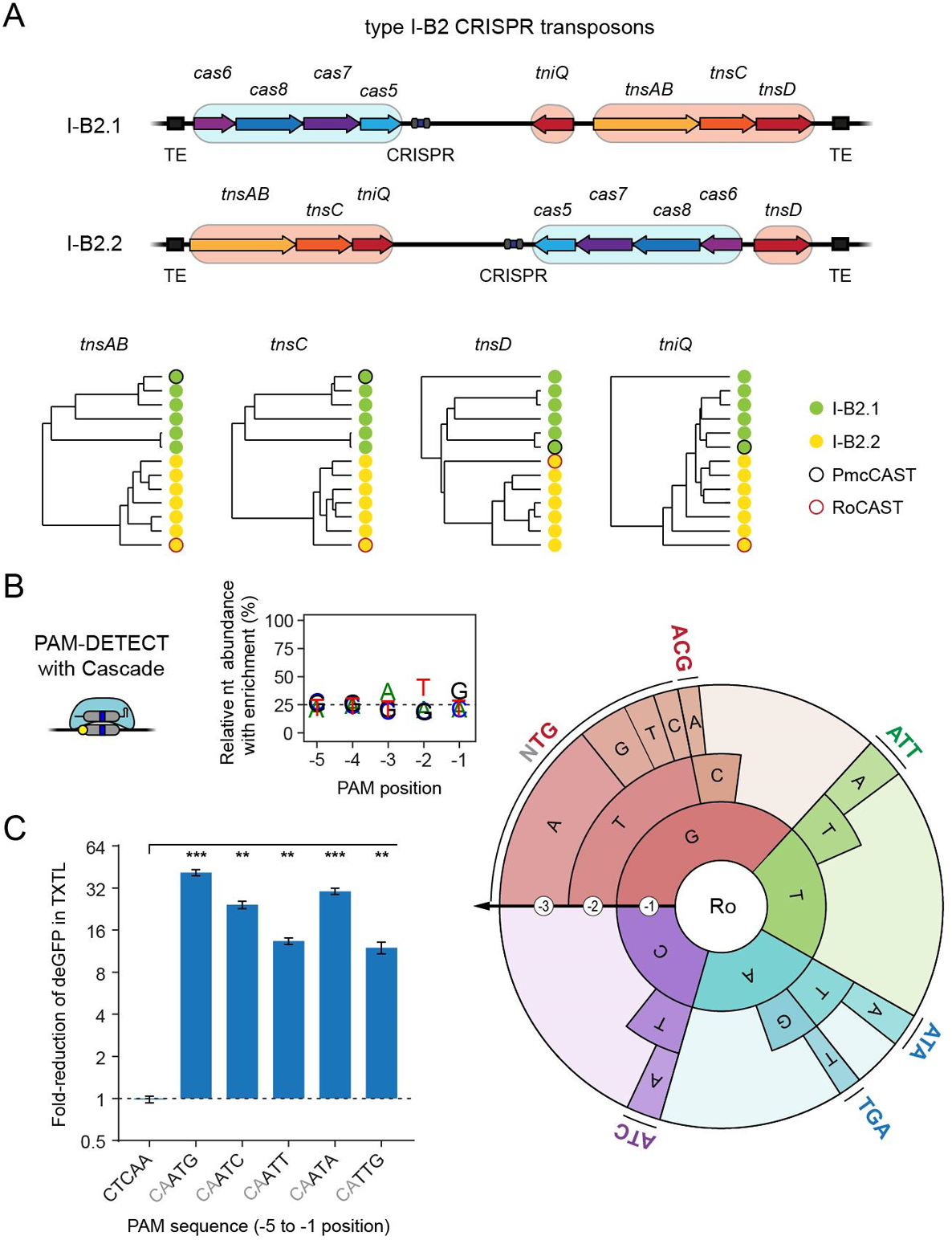
Interrogating PAM requirements of the *Rippkaea orientalis* I-B2.2 CRISPR transposon. (**A**) Overview of I-B2.1 and I-B2.2 CRISPR transposons. The two are divided based on the gene organization within each transposon. Phylogenetic trees are shown for the transposon genes. The *Peltigera membranacea* cyanobiont 210A CRISPR transposon (PmcCAST) from the I-B2.1 branch was previously characterized (Saito et al., 2021). The *R. orientalis* CRISPR transposon (RoCAST) from the I-B2.2 branch is characterized in this work. See Supplementary Figure S4 for alignments with names that match the order within the trees. (**B**) Nucleotide-enrichment plot and PAM wheel based on conducting PAM-DETECT with Cascade from RoCAST. Individual sequences comprising at least 2% of the PAM wheel are shown. The plot and PAM wheel are averages of duplicate independent experiments. (**C**) Validation of PAMs in TXTL. Gene repression was evaluated with Cascade and the indicated PAM flanking target R upstream of the deGFP reporter. See Figure 2E for details. Fold-reduction was calculated based on a non-targeting crRNA control. CTCAA matches the 3′ end of the repeat and therefore serves as a negative control. The CTCAA self PAM is the reference for statistical analyses. Error bars in C indicate the mean and standard deviation of triplicate independent experiments. ***: p < 0.001. **: p < 0.01. *: p < 0.05. ns: p > 0.05.

We conducted PAM-DETECT by expressing a single-spacer CRISPR array as well as the four RoCAST Cascade genes (*cas5*, *cas6*, *cas7*, *cas8*) from two separate expression constructs. This combination yielded a PAM profile dominated by ATG (**Figs. 6B, S4B and C**), matching the PAM recognized by the one previously characterized I-B2.1 CAST from *Peltigera membranacea cyanobiont* 210A (PmcCAST) (Saito et al., 2021). This match was expected given the high similarity (65-81%) between the protein components forming PmcCAST and RoCAST Cascade. However, single-nucleotide perturbations to ATG could be recognized by the RoCAST even under low Cascade conditions. The TXTL-based deGFP silencing assay confirmed recognition of ATG as well as the single-nucleotide perturbations (**Fig. 6C**). We further showed that PAM-DETECT can be applied to the previously characterized I-B1 CRISPR transposon from *Anabaena variabilis* ATCC 29413 (AvCAST) (Saito et al., 2021) (**Fig. S5A and B**). These insights came from using a streamlined TXTL assay without any protein or RNA purification and only half of the genetic components needed for transposition.

### DNA transposition by CRISPR transposons can be recapitulated in TXTL

We next wanted to evaluate how insights into PAM recognition translate into DNA transposition. However, doing so with *in vitro* or cell-based assays posed numerous challenges that would slow the characterization process. In particular, encoding and expressing all of the genetic components into a few compatible plasmids is laborious and could require extensive optimization, while overexpressing some components could be toxic to the cells. Instead, we asked whether transposition could be recapitulated in TXTL (**Fig. 7A**) to rapidly test different configurations and constructs.

**Figure 7.**
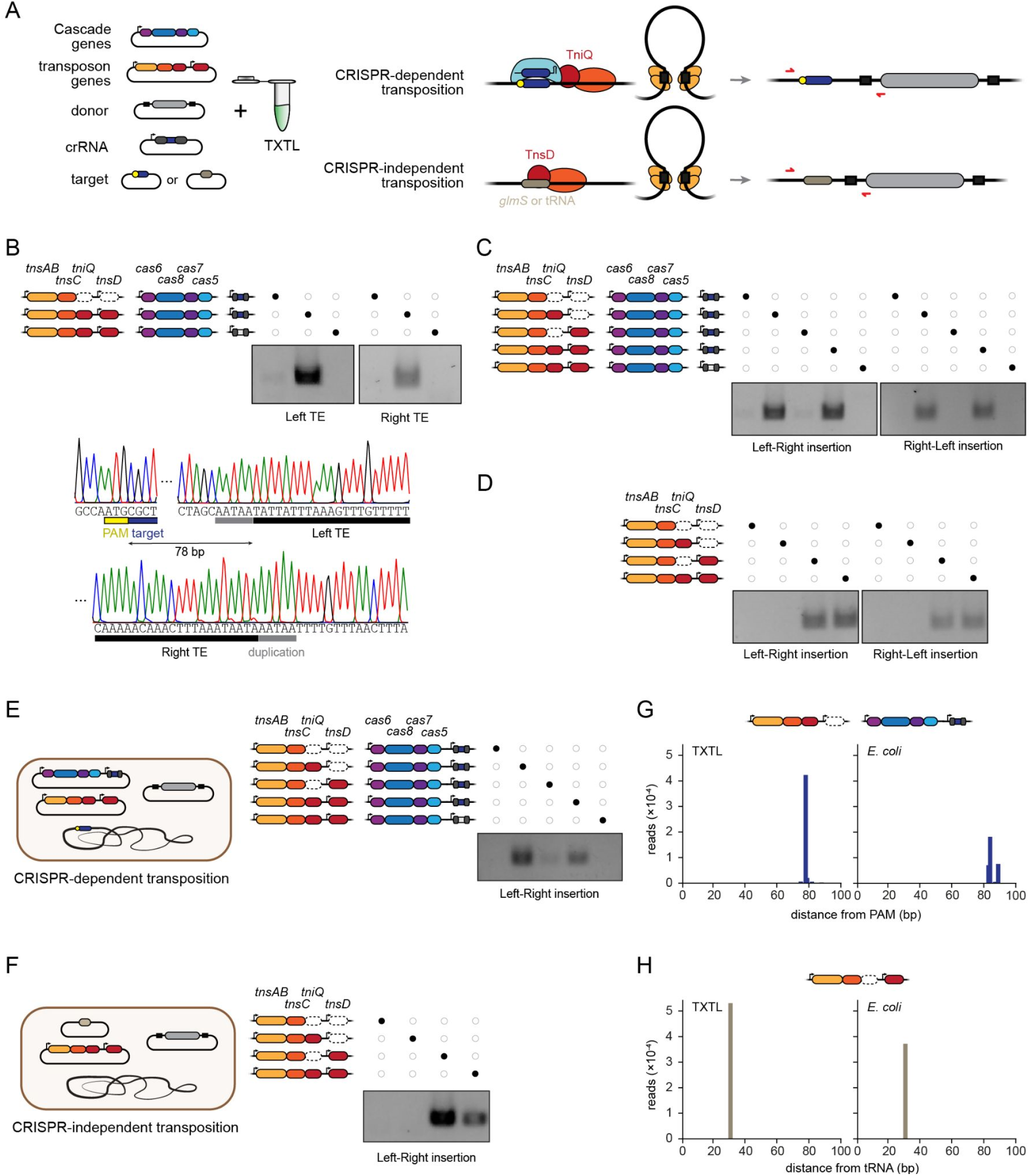
Investigating transposition of the *Rippkaea orientalis* I-B2.2 CRISPR transposon in TXTL and in *E. coli*. (**A**) Overview of the TXTL-based transposition assay. For I-B CRISPR transposons, transposition can occur through crRNA-guided recognition of a DNA target or through TnsD-guided recognition of *glmS* or a tRNA gene independent of the CRISPR machinery. Primers (red) are shown to selectively amplify the transposition product. The *R. orientalis* I-B2.2 CRISPR transposon (RoCAST) flanks the tRNA-Leu gene. (**B**) CRISPR-dependent transposition and determination of transposon ends and insertion distance using the TXTL-based transposition assay with RoCAST. PCR products are specific to the left-right orientation and span the crRNA target site and the beginning of the cargo (left TE) or the end of the cargo and downstream of the insertion site (right TE). (**C**) CRISPR-dependent transposition in TXTL. PCR products span the crRNA target site and the beginning of the cargo for both orientations of transposon insertion. (**D**) CRISPR-independent transposition in TXTL. PCR products span the end of the tRNA-Leu gene and the beginning of the cargo for both orientations of transposon insertion. (**E**) CRISPR-dependent transposition in *E. coli*. PCR products span the crRNA target site and the beginning of the cargo (left-right orientation). (**F**) CRISPR-independent transposition in *E. coli*. PCR products span the TnsD target site and the beginning of the cargo (left-right orientation). (**G**) Assessment of insertion distances for CRISPR-dependent transposition in TXTL and in *E. coli*. The constructs lacking *tnsD* were used. Transposition was determined by next-generation sequencing of the PCR product spanning the crRNA target site and the beginning of the cargo (left-right orientation). (**H**) Assessment of insertion distances for CRISPR-independent transposition in TXTL and in *E. coli*. The constructs lacking *tniQ* were used. Transposition was determined by next-generation sequencing of the PCR product spanning the end of the tRNA-Leu gene and the beginning of the cargo (left-right orientation). All gel images are representative of at least duplicate independent experiments.

We began with the *V. cholerae* I-F CAST. Combining DNA constructs encoding a targeting single-spacer array, three Cascade genes, four transposon genes (*tnsA*, *tnsB*, *tnsC*, *tniQ*), donor DNA flanked by the transposon ends, and a target construct resulted in measurable DNA transposition in both orientations by PCR (**Fig. S6A**). Sanger sequencing of the PCR products revealed the core transposon ends as well as the distance between the target site and insertion site that aligned with prior work (**Fig. S6A**). We were also able to reconstitute transposition in TXTL for AvCAST (**Fig. S5C**). Therefore, TXTL can be used to characterize DNA transposition by CASTs.

### DNA transposition in TXTL with the *Rippkaea orientalis* CAST establishes a distinct branch within I-B2 CRISPR transposons

Building on TXTL-based transposition with the I-F and I-B2.1 CASTs, we evaluated DNA transposition in TXTL with the I-B2.2 RoCAST (**Fig. 7B**). Because the ends of this transposon were unclear, we constructed a donor DNA construct flanked by two 250-bp sequences predicted to contain the right and left RoCAST ends. We combined the donor DNA and target DNA flanked by an ATG PAM with constructs encoding the I-B2.2 Cascade genes (*cas5*, *cas6*, *cas7*, *cas8*), transposase genes (*tnsAB, tnsC, tnsD, tniQ*), and a single-spacer CRISPR array with a targeting or non-targeting spacer. The TXTL reactions resulted in measurable crRNA-directed transposition in both orientations by PCR. Sanger sequencing of the PCR products revealed the core transposon ends along with five bases that are duplicated as part of transposition **(Fig. 7B)**, similar to other CASTs (Klompe et al., 2019).

Recent work revealed that I-B CASTs possess two distinct modes of transposition: CRISPR-dependent transposition through TniQ and DNA targeting by Cascade and CRISPR-independent transposition through TnsD (Saito et al., 2021). We therefore evaluated the role of TniQ and TnsD for either mode of transposition in TXTL. For CRISPR-dependent transposition, TXTL reactions with TniQ yielded the highest CRISPR-dependent transposition efficiency. However, we surprisingly observed modest but detectable crRNA-dependent transposition even in the absence of TniQ and TnsD by PCR (**Fig. 7B and C**) and by next-generation sequencing of the PCR product (**Fig. S6B**). As further support for I-B2.2 as a separate branch, TniQ was reported to be required for crRNA-dependent transposition by the I-B1 AvCAST (**Fig. S5C**) and the I-B2.1 PmcCAST (Saito et al., 2021). To explore CRISPR-independent transposition, we swapped the crRNA target for the tRNA-Leu gene naturally flanking RoCAST in the *R. orientalis* genome. CRISPR-independent transposition was detected in both orientations (**Fig. 7D**). Transposition required TnsAB, TnsC and TnsD, while removing TnsD or replacing it with TniQ eliminated transposition.

We finally asked how the properties of RoCAST observed in TXTL translate *in vivo*. We adapted the DNA constructs for use in *E. coli* by condensing the constructs into three plasmids (**Fig. 7E and F**). For CRISPR-dependent transposition, we targeted the *lacZ* gene in the *E. coli* genome at a site flanked by an ATG PAM. Over-expressing Cascade proved to be cytotoxic, reflecting challenges to characterizing CASTs *in vivo*, although the cytotoxicity could be relieved with minimal induction of Cascade expression. In line with the TXTL results, CRISPR-dependent transposition was measurable by PCR in *E. coli* strains expressing the Cascade, TnsAB, TnsC and TniQ proteins, albeit only for the left-to-right insertion orientation (**Fig. 7E**). Removing TnsD boosted this mode of transposition (**Fig. 7E**). Somewhat paralleling the TXTL results, less efficient transposition was measurable by PCR in the absence of TniQ but not both TniQ and TnsD (**Figs. 7E and S6C**). For CRISPR-independent transposition, we targeted a vector carrying the terminal region of the tRNA-Leu gene from the *R. orientalis* genome. Matching the TXTL results, TnsAB, TnsC, and TnsD proteins were necessary for transposition (**Fig. 7F**). To compare the insertion distances between the target and the inserted donor DNA in TXTL and in *E. coli*, the PCR products were subjected to next-generation sequencing. For CRISPR-dependent transposition, transposition in TXTL consistently occurred 78 bps downstream of the PAM, while transposition in *E. coli* principally occurred within a window of 83-89 bps downstream of the PAM (**Fig. 7G**), although the difference may be attributed to the use of different target sites and insertion contexts as was previously reported for the I-B1 AvCAST (Saito et al., 2021). For CRISPR-independent transposition, transposition in TXTL and in *E. coli* both occurred 31 bps downstream of the *tRNA-Leu* gene (**Fig. 7H**). The insertion distances for both modes of transposition are comparable to the insertion windows identified for the other characterized I-B2 system (Saito et al., 2021). Overall, these findings demonstrate that insights from TXTL-based transposition translate into *in vivo* settings.

## DISCUSSION

Through multiple demonstrations, we showed how cell-free TXTL reactions could be applied to rapidly characterize multi-component CRISPR nucleases as well as CRISPR transposons. One method we used repeatedly, PAM-DETECT, could comprehensively determine PAM sequences recognized by the DNA-binding machinery of an immune system or transposon. Our method offered important advantages over current cell-based and *in vitro*-based methods that should accelerate characterization of Class 1 CRISPR-Cas systems and transposons. PAM-DETECT could be completed in under one day starting from purified DNA constructs and ending with amplicons for next-generation sequencing. In contrast, cell-based methods require DNA transformation, culturing, and growth before DNA isolation that can stretch for days. *In vitro* assays can require even more time due to the need to purify ribonucleoprotein complexes overexpressed in cells. Both traditional methods can require extensive optimization, such as combining the constructs into a small set of compatible plasmids with appropriate expression, tackling issues of toxicity, or troubleshooting issues that arise during purification--steps that are irrelevant for TXTL. Finally, the ability to conduct reactions in a few microliters allows PAM-DETECT to be readily scaled, allowing the parallel interrogation of tens or even hundreds of systems under different reaction conditions. While TXTL reactions are normally conducted between 25°C and 37°C, the DNA-binding and restriction steps could be conducted at elevated temperatures, such as for evaluating CRISPR-Cas systems derived from thermophiles and hyperthermophiles. In addition, while overexpression of Cascade could lead to unwanted enrichment of suboptimal PAMs, we demonstrated how the reaction conditions could be tuned and how qPCR could be applied to gauge the extent of library protection. Given these advantages, TXTL-based characterization of Class 1 systems could represent a widespread means to explore these abundant and diverse systems.

We further leveraged TXTL to accelerate the validation and extension of our results from PAM-DETECT. We frequently employed a deGFP repression assay in which target binding by Cascade blocks deGFP expression. This assay allowed us to confirm PAM sequences, where deGFP repression strongly correlated with enrichment with PAM-DETECT for the *E. coli* I-E system. One potential limitation to PAM-DETECT and the repression assay is that binding may not correspond to DNA degradation, as was reported to some degree for DNA binding and degradation by the I-E system (Xue et al., 2015). However, as part of characterizing the self-targeting CRISPR-Cas systems in *X. albilineans*, we showed that the repression assay could be readily modified to specifically assess DNA degradation by Cas3. By targeting a location well upstream of the promoter, a reduction of deGFP expression would only occur through the action of Cas3. This altered setup could be readily applied to validate identified PAMs in the context of DNA degradation. Finally, we showed that DNA transposition by CASTs could be fully recapitulated in TXTL. We were able to recapitulate CRISPR-dependent and CRISPR-independent transposition by I-B and I-F CASTs, suggesting that TXTL would be valid for V-K CASTs representing the third and final subtype (Saito et al., 2021; Strecker et al., 2019). With these additional assays in place, TXTL can be applied well beyond PAM determination.

One major application we pursued was mining the natural diversity of I-E CRISPR-Cas systems. Using PAM-DETECT, we evaluated 11 different systems representing diverse sequences within the variable L1 loop of the Cas8e protein. The analysis revealed ranging extents of library protection indicative of Cascade expression, binding activity, and the breadth of recognized PAMs. The identified PAMs deviated from that associated with *E. coli*’s I-E system, suggesting that a far broader range of PAMs could be revealed by further interrogating the diversity of these systems. Whether the diversity parallels that observed for Cas9 nucleases remains to be seen and could reflect the distinct forces that shaped the evolution of each system type (Gasiunas et al., 2020). A similar approach could be particularly powerful for mining I-C and I-Fv Cascade complexes that require the fewest number of Cas proteins (Hochstrasser et al., 2016; Pausch et al., 2017). Complexes could be mined exhibiting not only unique PAM preferences but also smaller proteins, altered temperature ranges, or enhanced binding and cleavage activities. Given the proliferation of engineered single-effectors with altered PAM recognition (Collias and Beisel, 2021), TXTL could be applied to characterize any similarly engineered variants of type I systems.

Beyond mining orthologs within a CRISPR-Cas subtype, PAM-DETECT offered a powerful means to interrogate CRISPR-Cas systems with potentially unique properties. We specifically focused on a I-C system and a I-F1 system present in *X. albilineans* that encode a large repertoire of self-targeting spacers. While genetic deactivation of the CRISPR machinery is thought to be a common means of resolving otherwise lethal self-targeting (Stern et al., 2010), we showed that Cascade and Cas3 were functionally encoded and could recognize PAMs flanking the vast majority of the self targets. These findings instead suggest that the expression or activity of the CRISPR machinery is inhibited, preventing lethal self-targeting. One possibility is that the cell encodes Acrs that actively inhibit steps of CRISPR-based immunity or expression (Davidson et al., 2020). Future work therefore could interrogate what is preventing both systems from lethal self-targeting not only in *X. albilineans* but also the many other organisms possessing CRISPR-Cas systems with self-targeting spacers. This work could reveal novel classes of Acrs as well as instances of CRISPR-Cas systems performing functions extending beyond adaptive immunity.

As a final example, we applied TXTL to characterize a distinct branch of I-B2 CASTs. The I-B CAST type was recently divided into two subtypes (I-B1 and I-B2) based on whether *tnsA* and *tnsB* were fused, the genetic organization of the CAST, and the site recognized for CRISPR-independent insertion (Saito et al., 2021). When exploring I-B2 CASTs, we noticed a clear division in the genetic organization of these CASTs that paralleled phylogenetic trees for the transposon genes. We further found that CRISPR-dependent transposition could occur in the absence of TniQ for one branch (I-B2.2), contrasting with the essential role of TniQ described for the other branch (I-B2.1) and subtype (I-B1) (Saito et al., 2021). TniQ-independent transposition under these conditions was weak, raising questions whether CRISPR-dependent transposition would occur in the absence of TniQ under natural settings. Regardless of the biological relevance, it likely reflects distinct biomolecular mechanisms and interactions that further support some division in categorization. As only a small number of CASTs have been characterized to-date, further exploring these unique mobile genetic elements could reveal new properties and provide CASTs for further technological development and application. In that regard, applying cell-free systems could greatly aid these efforts and help drive new discoveries and technologies.

## ACKNOWLEDGEMENTS

We thank Natalia Ivanova for assistance with the bioinformatics identification of Type I-E CRISPR-Cas systems. We thank Sam Sternberg for providing plasmid pSL0283, pSL0284 and pSL0527. This work was supported by an ERC Consolidator grant (865973 to C.L.B.), the Deutsche Forschungsgemeinschaft (BE 6703/1-1 to C.L.B.), and the Netherlands Organization for Scientific Research (NWO) through a Rubicon Grant (project 019.193EN.032 to I.M.). A portion of this research was performed under the JGI-EMSL Collaborative Science Initiative and used resources at the DOE Joint Genome Institute and the Environmental Molecular Science Laboratory, which are DOE Office of Science User Facilities. Both facilities are sponsored by the Office of Biological and Environmental Research and operated under Contract Nos. DE-AC02-05CH11231 (JGI) and DE-AC05-76RL01830 (EMSL).

## AUTHOR CONTRIBUTIONS

Conceptualization: F.W., I.M., C.L.B.; Methodology: F.W., I.M., C.L.B., Software: F.W., I.M., Validation: F.W., I.M., F.E.; Investigation: F.W., I.M., F.E., Writing - Original Draft: F.W., I.M., C.L.B., Writing - Review & Editing: F.W., I.M., F.E., C.L.B. Visualization: F.W., I.M., C.L.B., Supervision: C.L.B.; Funding acquisition: C.L.B.

## DECLARATION OF INTERESTS

C.L.B. is a co-founder and member of the Scientific Advisory Board member for Locus Biosciences and is a member of the Scientific Advisory Board for Benson Hill. The other authors declare no competing interests.

## STAR METHODS

## METHOD DETAILS

### Plasmid construction

Standard cloning methods Gibson Assembly, Site Directed Mutagenesis (SDM) and Golden Gate were used to clone plasmids used in TXTL experiments. pPAM_library containing a PAM library with five randomized nucleotides was generated by SDM on p70a-deGFP_PacI with primers FW531 and FW532 (**Table S5**). Single-spacer CRISPR arrays were generated either with Golden Gate adding spacer sequences in a plasmid containing two repeat sequences interspaced by two BaeI or BbsI restriction sites or by SDM on pEc_gRNA1, pEc_gRNA2 or pEc_gRNAnt to change the repeat sequences to match the tested CRISPR systems. Plasmids harboring different PAM sequences for PAM validation assays were generated by SDM on p70a-deGFP_PacI. To generate plasmids encoding *X. albilineans* type I-C and type I-F1 Cas proteins, genomic DNA isolated from *Xanthomonas albilineans* CFBP7063 was PCR amplified using Q5 Hot Start High-Fidelity 2X Master Mix (NEB) and cloned into pET28a using Gibson Assembly. All other plasmids were generated with Gibson Assembly or SDM (**Table S5**). All constructed plasmids were verified with Sanger sequencing.

For the VcCAST *in vivo* transposition experiments we cloned into the previously described pSL0284 vector (Klompe et al., 2019) two spacers targeting the *lacZ* gene of the *E. coli* BL21 (DE3) genome, yielding the pQCas_CAA and pQCas_AAA vectors. The protospacer targeted by the former vector has a 5’CAA PAM, whereas the protospacer targeted by the latter vector has a 5’AAA PAM.

For the RoCAST *in vivo* transposition experiments, genes encoding the *Rippkaea orientalis tnsAB, tnsC, tnsD and tniQ* were synthesized (Twist Bioscience) and cloned in the pET24a vector in various combinations, resulting in the construction of the pRoTnsABC, pRoTnsABCD, pRoTnsABCQ, pRoTnsABCDQ vectors (**Table S5**). The *Rippkaea orientalis* Cascade operon (*cas6, cas8, cas7, cas5*) was synthesized (Twist Bioscience) and cloned into the pCDFDuet-1 vector together with a *gfp* gene flanked by two BsaI restriction sites and the corresponding CRISPR direct repeats. Into the resulting pRoCascade_gfp vector we cloned a spacer targeting the *lacZ* gene of the *E. coli* BL21 (DE3) genome and a non-targeting control spacer, constructing the pRoCascade_T (targeting) and pRoCascade_NT (non-targeting) vectors, respectively (**Table S5**). DNA fragments encoding the right and left RoCAST ends were synthesized (IDT) and cloned into the pUC19 vector flanking a *gfp* gene, yielding pRoDonor (**Table S5**). A 105-bp long DNA fragment from the *Rippkaea orientalis* genome, encoding the region which is located right upstream of the left end of RoCAST and includes the last 74 bp of the *tRNA-Leu* gene, was synthesized (IDT) and cloned into the pCDFDuet-1 vector, resulting in the construction of the pRoTarget vector (**Table S5**).

## PAM-DETECT

A plasmid with five randomized nucleotides flanking a target site covering a PacI restriction enzyme recognition site was constructed as described before. If Cas proteins required for Cascade formation were encoded on separate plasmids, a MasterMix with the required Cas protein encoding plasmids in their stoichiometric amount was prepared beforehand. Thereby, a stoichiometry of Cas8e_1_-Cse2_2_-Cas_76_-Cas5_1_-Cas6_1_ was used for all Type I-E systems. A 6 µL TXTL reaction was assembled consisting of 3 nM (high Cascade) or 0.25 nM (low Cascade) of the Cascade-encoding plasmid or the Cascade MasterMix, 4.5 µL myTXTL Sigma 70 Master Mix, 0.2 nM pET28a_T7RNAP, 0.5 mM IPTG, 1 nM gRNA-encoding plasmid and 1 nM pPAM_library. A negative control containing all components from the reaction besides the Cascade plasmids and the gRNA-expressing plasmid was included. PAM-DETECT assays assessing either the type I-C or the type I-F1 system in *X. albilineans* were lacking IPTG in their reactions. TXTL reactions were incubated at 29°C for 6 h or 16 h. The samples were diluted 1:400 in nuclease-free H2O. 500 µL were digested at 37°C with PacI (NEB) at 0.09 units/µL in 1x CutSmart Buffer (NEB) for 1 h and 500 µL were used as a “non-digested” control by adding nuclease-free H2O instead of PacI. After inactivation of PacI at 65°C for 20 min, 0.05 mg/mL Proteinase K (GE Healthcare) was added and incubated at 45°C for 1 h. After inactivation of Proteinase K at 95 °C for 5 min, remaining plasmids were extracted via standard EtOH precipitation. Illumina adapters with unique dual indices were added by two amplification steps with KAPA HiFi HotStart Library Amplification Kit (KAPA Biosystems) and purified by Agencourt AMPure XP (Beckman Coulter) after every PCR reaction. The first PCR reaction adds the Illumina sequencing primers with primers that can be found in Table S5 using 15 µL of the EtOH-purified samples in a 50 µL reaction and 19 cycles. The second PCR adds the unique dual indices and the flow cell binding sequence using 1 ng purified amplicons generated with the first PCR using 18 cycles. The samples were submitted for next-generation sequencing with 50 bp paired-end reads with 1.25 or 2.0 million reads per sample on an Illumina NovaSeq 6000 sequencer. PAM wheels were generated according to Leenay et al. (Leenay et al., 2016). Nucleotide enrichment plot generation was adapted to the script from Marshall et al. (Marshall et al., 2018) by changing the script to visualize the probability of a given nucleotide at a given position by depicting the percentage of the nucleotide in that position. All PAM-DETECT assays were done in duplicates and PAM wheel and nucleotide enrichment plots show averages. The generated NGS data have been deposited in NCBI’s Gene Expression Omnibus (Edgar et al., 2002) and are accessible through GEO Series accession number GSE179614 (https://www.ncbi.nlm.nih.gov/geo/query/acc.cgi?acc=GSE179614 ). The following token can be used to access the data prior to publication: exexiqgyhrcblgj.

### qPCR Reactions

To assess the remaining amount of PAM-library containing plasmid after conducting PAM-DETECT, quantitative PCR (qPCR) was performed using SsoAdvanced Universal SYBR Green Supermix (Biorad) in 10 µL reactions. The reactions were quantified using a QuantStudio Real-Time PCR System (Thermo Fisher) with an annealing temperature of 68 °C according to manufacturers’ instructions. All samples were prepared by using the liquid handling machine Echo525 (Beckman Coulter).

### deGFP repression assays in TXTL

To assess activity of CRISPR-Cas systems, deGFP-repression assays in 3 µL or 5 µL TXTL reactions were conducted, measuring deGFP-expression over time in a 96-well V-bottom plate with BioTek Synergy H1 plate reader (BioTek) at 485/528 nm excitation/emission (Shin and Noireaux, 2012). All TXTL samples were either prepared by hand or by using the liquid handling machine Echo525 (Beckman Coulter).

3 µL TXTL reactions for PAM validation assays were prepared containing Cascade plasmid concentrations according to Table S2. If Cas proteins required for Cascade formation were encoded on separate plasmids, a MasterMix with the required Cas protein encoding plasmids in their stoichiometric amount was prepared beforehand. Thereby, a stoichiometry of Cas5_1_-Cas8_1_-Cas7_7_ was used for *X. albilineans* Type I-C, Cas8f1_1_-Cas5f1_1_-Cas7f1_6_-Cas6f_1_ was used for *X. albilineans* Type I-F1 and Cas8e_1_-Cse2_2_-Cas7_6_-Cas5_1_-Cas6_1_ was used for all Type I-E systems. Other components included in the TXTL reactions were 2.25 µL myTXTL Sigma 70 Master Mix, 0.2 nM p70a_T7RNAP, 0.5 mM IPTG and 1 nM gRNA-encoding plasmid. After a 4 h pre-incubation at 29 °C or 37 °C that allowed the ribonucleoprotein complex of Cascade and crRNA to form, 1 nM reporter plasmid (pGFP_XXXXX) with various PAM sequences in close proximity to the promoter driving deGFP expression was added to the reaction to ensure Cascade-binding would lead to deGFP inhibition. The reactions were incubated for additional 16 h at 29 °C or 37 °C while measuring deGFP expression. The gRNAs were constructed to target a protospacer within the *degfp* promoter located adjacent to the various PAM sequences.

To test the cleavage and/or binding ability of the type I-C and the type I-F1 systems in *X. albilineans*, 3 µL TXTL assays were conducted containing Cascade-encoding plasmids in the stoichiometry as mentioned before. To test binding ability, 2.25 µL myTXTL Sigma 70 Master Mix, 0.2 nM p70a_T7RNAP, 0.5 mM IPTG, 1 nM gRNA1-, gRNA2, or gRNAnt-encoding plasmid and 1 nM or 0.25 nM Cascade MasterMix was added to a TXTL reaction for the type I-C and type I-F1 system, respectively. To test cleavage ability, 2.25 µL myTXTL Sigma 70 Master Mix, 0.2 nM p70a_T7RNAP, 0.5 mM IPTG, 1 nM gRNA1-, gRNA2, or gRNAnt-encoding plasmid, 1 nM Cascade MasterMix and 0.5 nM or 0.25 nM pXalb_IC_Cas3 or pXalb_IF_Cas2-3 was added to a TXTL reaction for the type I-C and type I-F1 system, respectively. After 4 h pre-expression at 29°C, 1 nM p70a_deGFP reporter plasmid was added to the reactions and incubated for additional 16 h at 29°C while measuring deGFP-fluorescence. gRNA1 is designed to target a protospacer within the promoter driving deGFP expression adjacent to a type I-C TTC or a type I-F1 CC PAM to ensure Cascade-binding would lead to deGFP-inhibition. gRNA2 is designed to target a protospacer adjacent to a type I-C TTC or a type I-F1 CC PAM upstream of the promoter to ensure cleavage of the targeted plasmid would result in deGFP-inhibition whereas binding-only would result in deGFP-production. gRNAnt represents a non-targeting control.

5 µL TXTL reactions assessing dispensability of TniQ for *V. cholerae* I-F CAST Cascade-binding were performed with reactions containing 3.75 µL myTXTL Sigma 70 Master Mix, 0.2 nM p70a_T7RNAP, 0.5 mM IPTG and 0.5 nM pVch_IF_CasQ_gRNA3/nt or 0.5 nM pVch_IF_Cas_gRNA3/nt. After a 4 h pre-incubation step at 29 °C, the reporter plasmid p70a_deGFP was added and the reactions were incubated for additional 16 h at 29 °C while measuring deGFP-fluorescence. gRNA3 is designed to target a protospacer within the promoter driving deGFP expression adjacent to a CC PAM. gRNAnt represents a non-targeting control.

### Transposition in TXTL

To assess crRNA-dependent transposition of the *Vibrio cholerae Tn6677* I-F CAST in TXTL, 5 µL TXTL reactions containing 3.75 µL myTXTL Sigma 70 Master Mix, 0.2 nM p70a_T7RNAP, 0.5 mM IPTG, 1 nM of the previously described donor plasmid (pSL0527), 2 nM of the previously described TnsABC-plasmid (pSL0283) (Klompe et al., 2019), 1 nM p70a_deGFP and 1 nM pVch_IF_CasQ_gRNA3 or pVch_IF_CasQ_gRNAnt were prepared. The reactions were incubated at 29 °C for 16 h. Transposition events were detected in a 1:400 dilution of the TXTL reaction by PCR amplification using Q5 Hot Start High-Fidelity 2X Master Mix (NEB) and combinations of donor DNA and genome specific primers. Transposition was verified by Sanger sequencing (**Table S5**).

crRNA-dependent transposition of RoCAST in TXTL was performed in 3 µL TXTL reactions consisting of 2.25 µL myTXTL Sigma 70 Master Mix, 0.2 nM p70a_T7RNAP, 0.5 mM IPTG, 1 nM pRoCascade, 1 nM pRo_gRNA2/nt, 1 nM pGFP_CAATG, 1 nM pRoDonor or pRoDonor_extended and 1 nM pRoTnsABC, pRoTnsABCD, pRoTnsABCQ or pRoTnsABCDQ. The reactions were incubated at 29 °C for 16 h. Transposition events were detected in a 1:100 dilution of the TXTL reaction by PCR amplification using Q5 Hot Start High-Fidelity 2X Master Mix (NEB) and combinations of donor DNA and genome specific primers (**Table S5**). Transposition was verified by Sanger sequencing.

crRNA-independent transposition of RoCAST in TXTL was performed in 3 µL TXTL reactions consisting of 2.25 µL myTXTL Sigma 70 Master Mix, 0.2 nM p70a_T7RNAP, 0.5 mM IPTG, 1 nM pRoTarget, 1 nM pRoDonor and 1 nM pRoTnsABC, pRoTnsABCD, pRoTnsABCQ or pRoTnsABCDQ. The reactions were incubated at 29 °C for 16 h. Transposition events were detected in a 1:100 dilution of the TXTL reaction by PCR amplification using Q5 Hot Start High-Fidelity 2X Master Mix (NEB) and combinations of donor DNA and genome specific primers (**Table S5**). Transposition was verified by Sanger sequencing.

### Transposition *in vivo*

For the crRNA-dependent transposition *in vivo* using the I-F CAST from *Vibrio cholerae Tn6677*, we employed the previously described transposition system (Klompe et al., 2019). We electroporated 30 ng of the pSL0283 vector with 30 ng of the pSL0527 vector and 30 ng of either the pQCas_CAA or pQCas_AAA vector into *E. coli* BL21(DE3) electrocompetent cells. We plated a fraction of each electroporation mixture on 100 mg/ml ampicillin, 50 mg/ml spectinomycin, 50 mg/ml kanamycin, 0.1 mM IPTG and 100 µg/ml X-gal containing LB-agar plates. The plates were incubated for 24 h at 30°C and the formed colonies were subjected to blue/white screening. Transposition events were identified by colony PCR using Q5 Hot Start High-Fidelity 2X Master Mix (NEB) and genome specific primers (**Table S5**).

For the crRNA-dependent transposition *in vivo* using RoCAST, we electroporated 30 ng of either pRoCascade_T or pRoCascade_NT vector with 30 ng of pRoDonor and 30 ng of either pRoTnsABC, pRoTnsABCD, pRoTnsABCQ or pRoTnsABCDQ vector into *E. coli* BL21(DE3) electrocompetent cells. We plated a fraction of each electroporation mixture on 100 mg/ml ampicillin, 50 mg/ml spectinomycin, and 50 mg/ml kanamycin containing LB-agar plates. The plates were incubated for 20 h at 37°C and the formed colonies were scraped and resuspended in LB liquid medium. A fraction of each cell suspension was re-plated on LB-agar plates supplemented with 100 mg/ml ampicillin, 50 mg/ml spectinomycin, 50 mg/ml kanamycin and 0.01 mM IPTG for induction of the expression of the Cascade and transposase proteins. The plates were incubated 20 h at 37°C and all the formed colonies were scraped and resuspended in LB liquid medium. A fraction of each cell suspension was subjected to gDNA isolation using the illustra Bacteria genomicPrep Mini Spin Kit (GE Healthcare). Transposition events were identified by PCR using Q5 Hot Start High-Fidelity 2X Master Mix (NEB) and combinations of donor DNA and genome specific primers (**Table S5**).

For the crRNA-independent *in vivo* transposition using RoCAST, we electroporated 30 ng of the pRoTarget with 30 ng of pRoDonor and 30 ng of either the pRoTnsABC, pRoTnsABCD, pRoTnsABCQ or pRoTnsABCDQ vector into *E. coli* BL21(DE3) electrocompetent cells. We plated a fraction of each electroporation mixture on 100 mg/ml ampicillin, 50 mg/ml spectinomycin, and 50 mg/ml kanamycin containing LB-agar plates. The plates were incubated for 20 h at 37°C and the formed colonies were scraped and resuspended in LB liquid medium. A fraction of each cell suspension was re-plated on LB-agar plates supplemented with 100 mg/ml ampicillin, 50 mg/ml spectinomycin, 50 mg/ml kanamycin and 0.01 mM IPTG for induction of the expression of the transposase proteins . The plates were incubated 20 h at 37°C and all the formed colonies were scraped and resuspended in LB liquid medium. A fraction of each cell suspension was subjected to gDNA isolation using the illustra Bacteria genomicPrep Mini Spin Kit (GE Healthcare). Transposition events were identified by PCR using Q5 Hot Start High-Fidelity 2X Master Mix (NEB) and combinations of donor DNA and pRoTarget specific primers (**Table S5**).

### Assessing transposition insertion point

To assess the exact insertion point of *Rippkaea orientalis* I-B2.2 CAST, *in vivo* and *in vitro,* transposition assays were conducted as previously described and the transposition products were PCR amplified and sent for next-generation sequencing. Illumina adapters with unique dual indices were added by two amplification steps with KAPA HiFi HotStart Library Amplification Kit (KAPA Biosystems) and each amplicon was purified by Agencourt AMPure XP (Beckman Coulter). The first PCR reaction adds the Illumina sequencing primer sites with primers that can be found in Table S5, the second PCR adds the unique dual indices and the flow cell binding sequences. 2 µL of 1:100 dilutions were used in a 50 µL PCR reaction to amplify TXTL reactions using either 19 or 30 cycles. 50 ng of genomic DNA were used in a 50 µL PCR reaction to amplify *in vivo* transposition with either 19 or 30 cycles. 1 ng of purified TXTL or *in vivo*-amplicon were subjected to the second PCR using 18 cycles. Library-pools consisting of six samples were submitted for next-generation sequencing with 300 paired-end reads with 0.15 million reads on an Illumina MiSeq machine.

The generated NGS data have been deposited in NCBI’s Gene Expression Omnibus (Edgar et al., 2002) and are accessible through GEO Series accession number GSE179614 (https://www.ncbi.nlm.nih.gov/geo/query/acc.cgi?acc=GSE179614). The following token can be used to access the data prior to publication: exexiqgyhrcblgj.

## QUANTIFICATION AND STATISTICAL ANALYSIS

### deGFP repression assays in TXTL

The fluorescence background was subtracted from the endpoint deGFP values with TXTL samples consisting of only myTXTL Sigma 70 Master Mix and nuclease-free water. The resulting endpoint deGFP values were either depicted as averages of a targeting gRNA and a non-targeting gRNA or fold change-repression was calculated by the ratio of non-targeting over the targeting deGFP values. Significance was calculated with Welch’s t-test. P > 0.05 is shown as ns, P < 0.05 is shown as *, P < 0.01 is shown as ** and P < 0.001 is shown as ***. Within the PAM validation assays represented as fold changes, significance was calculated between the fold change of a given PAM and the fold change of a PAM that corresponds to the 3’ end of the repeat of the tested CRISPR system. The fold changes of the PAM validation in Fig. 3B are depicted in a heat map. Thereby a difference between a non-targeting sample and a targeting sample with a specific PAM resulting in P > 0.05 is shown in white and excluded from further analysis. For all other samples within the heat map, the fold changes were calculated as mentioned above and presented relative to the highest fold change within one system. Significance within the deGFP repression assays testing binding and cleavage ability of the type I-C and the type I-F1 system in *X. albilineans* was calculated with the targeting and non-targeting sample for each condition. For the endpoint measurements in Fig. 5C, significance was calculated between a non-targeting sample and a targeting sample targeting the same PAM.

### qPCR

Cq values were used to measure target amounts. To calculate the relative abundance of the PAM library containing plasmid in the digested sample to the non-digested sample, the relative plasmid amount was normalized to a control amplifying the pET28a-T7RNAP that has no PacI recognition site using the the 2^(-(ddCt) method. Significance to the control sample lacking a CRISPR-Cas system was calculated with Welch’s t-test. P > 0.05 is shown as ns, P < 0.05 is shown as *, P < 0.01 is shown as ** and P < 0.001 is shown as *** .

### Assessing transposition insertion point

∼15 nts long sequences 5’ of the transposon terminal left end were extracted, counted and sorted. The sequences were mapped to the targeted plasmid or the targeted genome tolerating 2 nts mismatches and the distance between the insertion point and the PAM upstream of the protospacer or the end of the *tRNA-Leu* gene was noted. To only depict reliable insertion points, we present insertion points with more than 20 reads. The insertion points are shown as bar graphs.

The processed NGS data have been deposited in NCBI’s Gene Expression Omnibus (Edgar et al., 2002) and are accessible through GEO Series accession number GSE179614 (https://www.ncbi.nlm.nih.gov/geo/query/acc.cgi?acc=GSE179614). The following token can be used to access the data prior to publication: exexiqgyhrcblgj.

### *In silico* selection of representative type I-E CRISPR-Cas systems for PAM-DETECT

HMM profiles for the Cas5e, Cas6e, Cas7e and Cas8e proteins were developed upon aligning the members of the corresponding protein families (Cas5e: pfam09704, TIGR1868, TIGR02593; Cas6e: pfam08798, TIGR01907; Cas7e: pfam09344, TIGR01869; Cas8e: pfam 09481, TIGR02547). A new HMM profile was generated for the less conserved Cse2 protein upon aligning sequences with known 3D structure using PROMALS3D server (Pei et al., 2008) followed by a series of iterative alignment/model building steps to include additional sequences and increase sequence diversity. For the aligning processes of all five proteins, sequences were dereplicated at 90% identity using cd-hit (Huang et al., 2010) (with options -c 0.90 -g 1 -aS 0.9). The dereplicated sequences were compared against each other using blastp from blast+ v2.6.0 (Altschul et al., 1990) with e-value 10e-05 and defaults for the rest of parameters. Hits were filtered to retain those at >=60% pairwise identity, and were next clustered using the mcl algorithm (Enright et al., 2002) with inflation parameter of 2.0. Clusters with >=10 members were aligned using Gismo (Neuwald and Liu, 2004) with default parameters, and consensus sequences were extracted from the alignments. These consensus sequences, as well as singletons and sequences from smaller clusters were aligned using Gismo (Neuwald and Liu, 2004). Alignments were manually curated to remove shorter sequences that did not have one or more of the active site positions and HMM profiles were generated using hmmbuild (Eddy, 2009). Hmmsearch (Eddy, 2009) using the generated HMM profiles against all public genomes (isolates, SAGs, and MAGs), and all public metagenomes resulted in hits which were subsequently aligned against the generated HMM profiles. After selecting gene arrays that have all five complete or nearly complete genes, we identified 6,964 arrays in public genomes and 5,000 arrays in public metagenomes. Aligned sequences for all proteins from the same array were concatenated, and the resulting sequences were dereplicated with cd-hit (Huang et al., 2010) at 90% identity, aligned over at least 90% of the shorter sequences. This resulted in 2851 clusters, 1799 from metagenomes and 1052 from genomes. Whereas the alignment of the Cas8e proteins from these clusters showed high variability, the predicted L1 helix regions of the Cas8e, which have been shown to directly interact with the PAM (Xiao et al., 2017), presented higher conservation. We generated a list with the L1 signatures from the dereplicated cluster set and we subsequently manually filtered out systems that do not belong to known cultured mesophilic bacteria (**Table S3**). From the resulting list we selected I-E CRISPR/Cas systems with a variety of L1 motifs for experimental validation with PAM-DETECT.

### Comparative analysis of I-B CAST transposases

We searched previous literature (Peters et al., 2017; Saito et al., 2021) for *in silico* identified I-B2 CASTs, which contain a fused *tnsAB* gene and are easily distinguished from I-B1 CASTs, which contain separate *tnsA* and *tnsB* genes. We observed that one clade of the I-B2 CASTs encompasses systems with *tnsAB-tnsC-tnsD* operons while having the *tniQ* gene separated, whereas the other clade encompasses systems with *tnsAB-tnsC-tniQ* operons and the *tnsD* gene separated. We denoted the systems in the former clade as I-B2.1 CASTs and in the latter clade as I-B2.2 CASTs. We focused on the I-B2.2 CAST clade, that has no *in vitro* or *in vivo* characterized members, and we discarded from further analysis the systems that lacked at least one of the CRISPR-Cas or transposition genes (*tnsAB, tnsC, tnsD, tniQ, cas5, cas6, cas7, cas8*). We performed BlastP search (Altschul et al., 1990) using the TnsAB, TnsC, TnsD, TniQ proteins of each selected I-B2.2 system as queries, aiming to identify additional I-B2.2 CAST candidates. Our analysis yielded in total seven I-B2.2 systems and we selected six previously described I-B2.1 systems for phylogenetic analysis (Saito et al., 2021). The alignment of I-B2.1 and I-B2.2 transposition proteins was performed using T-Coffee (Di Tommaso et al., 2011), the phylogenetic trees were built using average distance and the BLOSUM62 matrix and they were visualized with JalView (Waterhouse et al., 2009).

### *In silico* analysis of RoCAST

We predicted the CRISPR array of RoCAST by uploading the *Rippkaea orientalis* genomic region between the *Rocas5* and *RotniQ* to CRISPRFinder (Grissa et al., 2007). The RoCAST ends were determined manually on Benchling by searching for repeat sequences of 20 nucleotides, with maximum 5 mismatched nucleotides, within the *Rippkaea orientalis* genomic regions 1 kb upstream of the *R. orientalis tnsAB* and 1 kb downstream of the *RotnsD.* We identified two types of repeat sequences present in both regions in opposite orientations and a candidate duplication region. Notably, we identified five repeat sequences in the predicted left end region, with one of the repeat sequences located downstream of the predicted duplication site, hence outside of the predicted RoCAST limits. The TXTL transposition demonstrated that this repeat is not part of the RoCAST transposon.

